# Simple droplet microfluidics platform for drug screening on cancer spheroids

**DOI:** 10.1101/2023.05.16.540960

**Authors:** Caroline Parent, Kiran Raj Melayil, Ya Zhou, Vivian Aubert, Didier Surdez, Olivier Delattre, Claire Wilhelm, Jean-Louis Viovy

## Abstract

3D in vitro biological systems are progressively replacing 2D systems to increase the physiological relevance of cellular studies. Microfluidics-based approaches can be powerful tools towards such biomimetic systems, but often require high-end complicated and expensive processes and equipments for microfabrication. Herein, a drug screening platform is proposed, minimizing technicality and manufacturing steps. It provides an alternate way of spheroid generation in droplets in tubes. Droplets microfluidics then elicit multiple droplets merging events at programmable times, to submit sequentially the spheroids to chemotherapy and to reagents for cytotoxicity screening. After a comprehensive study of tumorigenesis within the droplets, the system is validated for drug screening (IC50) with chemotherapies in cancer cell lines as well as cells from patient-derived-xenografts (PDX). As compared to microtiter plates methods, our sytem reduces the initial amout of cell up to 10 times and opens new avenues towards primary tumors drug screening approaches.

## Introduction

Thanks to the constant advances in biology and medicine, progresses in the understanding of diseases at the cellular and molecular levels occur rapidly. In cancer research, these progresses revealed the existence of heterogeneities, both intra and inter-tumoral, and their importance for therapeutic escape and treatment orientation.^1,2^ This knowledge is now paving the way for new therapeutic strategies. This triggered in particular the development of “precision medicine”, in which treatment are selected, within a constantly increasing drugs toolbox, based on patient-specific molecular biomarkers.^3^ This approach has provided considerable successes, but unfortunately a strong variability among patients remains in treatment responses, and numerous therapeutic escapes and/or treatment failures are still observed.^4–7^ Indeed, the intra-tumoral heterogeneity can hide, for instance, minority cancer cell population insensitive to the drug(s) active on the dominant cancer cells, more complex multicellular synergies between different subpopulations, or possible redundant paths for therapeutic. To overcome this difficulty and progress in treatment orientation beyond biomarkers-based precision medicine, appeared the idea of a more heuristic approach, in which drugs and/or drug combinations are tested directly on primary patient cells.^8^ This requires, however, the availability of enough cells for such systematic screening, and of cost-affordable methodologies.

The conventional drug testing method is based on 2D cancer cells culture. The 2D format is easy to implement, including at high throughput thanks to microtiter plates (MTP). They represent a massively adopted standard in biology. MTP allow for the implementation of relatively low-cost screening thanks to robots, and for easy high-resolution imaging. However, the MTP format requires a large number of cells, and poorly reproduces the actual metabolism of tumours.^9^ Indeed, in an organism, cells are most often organized in 3D structures, and this is also the case for solid tumours. This organization induces specific properties of cell-cell interactions and species transport (from oxygen to more complex signalling molecules),^10^ which cannot be adequately modelled by 2D cultures. The culture of cancer cells as 3D spheroids, coined in this case “tumoroids”, can overcome at least to some extent the limitations of 2D cultures mentioned above.^9,11^

The current formats for the growth and study of 3D cellular spheroids, based on non-adherent microtiter plates, also present a number of disadvantages. The presence of a rigid bottom surface, even a non-adhesive one, creates a strong and non-physiological mechanical constraint to the growth of cells, and biases asymmetrically the transport of nutriments and signalling molecules. This limitation can be overcome to some extent by the hanging drop technology, in which the spheroid grows without touching a solid surface, but it requires more complex manipulation and a higher cost. Also, in hanging drop method, imaging, buffer exchange and cell recovery are more delicate than in conventional MTP.^12,13^ Another technique, consisting in growing spheroids in 3D in hydrogels, creates a more physiological environment and strongly reduces the above problems of asymmetry, but it introduces other difficulties. Because of the random position of spheroids in the gel, access to nutrients by diffusion is uneven, the grown spheroids are generally very heterogeneous in sizes, and finally, imaging at high resolution and recovery of individual spheroids are problematic.^14^

Finally, all these methods typically require several thousand to several hundreds of thousands of cells per well. Besides, with the current relevant panel of possible treatments for each cancer type (and the general trend of combining the compounds), drug testing involves tens of different treatment formulations for each patient. Put together, this is incompatible with the number of cells available with the vast majority of current biopsy methods. Indeed, the development of minimally invasive biopsies such as Fine Needle Aspiration that harvests dozens to hundreds of thousands of cells and the (fortunate) evolution of cancer diagnosis towards earlier detection tend to reduce the size of patient cell samples.^15^ Reaching the number of cells required for conventional multicompound drug testing thus requires amplification steps, either by cell culture (which generally induces problematic phenotypic drift),^16,17^ or by a lengthy and costly implementation of several intermediate steps of multiplication of the initial sample in animal models. In this approach, called PDX for “Patients Derived Xenografts”, particularly developed in cancer research, tumour cells from a patient are injected in immune-supressed mice, leading to a “human tumour” grown to a size about 1cm (limited by ethical regulations), collected after mice euthanasia, and in general dissociated and re-injected into several generations of mice until sufficient material is obtained. This is thus a very expensive and time-consuming process (typically several months), ruining most of the initial simplicity and ethical benefit of the spheroids format. For instance, a typical large-scale screening of anticancer drugs requires several large robots, 9700 MTP and hundreds of mice. Finally, even if PDX yields less biological drift than multiple passes of in vitro culture, a drift from the phenotype of the initial cells is always possible, and difficult to assess.^18^

Thanks to their predisposition for miniaturization, microfluidic approaches appear as natural candidates for tumoroid culture to perform drug screening on small number of cells.^8,19^ Microfluidic platforms for all-in-one tumoroids culture, drug testing and results analysis present a promising innovation in personalized medicine for fast and low-cost drug testing. Indeed, some interesting platforms along this line have been proposed (Merten *et al*.,^20^ Tomasi *et al*.^21^) (we shall come back to these solutions in the discussion section).

In this project, we encapsulate cells in “plugs” (i.e. elongated droplets suspended in a carrier oil and highly confined in a capillary), and cultivate them until the formation of an individual tumoroid inside each plug. The velocity of the droplets in the microchannel depends on the relative viscosity of the fluids and of the droplet size.^22,23^ More precisely, at a fast enough flow rate, small droplets are faster than bigger ones, and the velocity difference is nonlinear versus flow rate. Using these properties, we can merge on-demand droplets containing drugs and droplets containing tumoroids to assess the drug efficiency. Our protocol involves “trains” of plugs containing either samples or reagents, which can be further mixed when desired. This feature is new and unique in microfluidic cell culture platforms, and in droplet microfluidics. It allows the implementation of complex, multi-steps protocols. This format also provides very reproducible tumoroids and allows high resolution imaging.

We demonstrate the possibility of growing tumoroids in droplets for 3-4 days, starting from about 350 cells in droplets of 700nL. We performed drug screening using a metabolic assay, which does not hypothesize any specific mechanism of action of the drug. It was applied to tumoroids formed from two different cancer cell lines and to tumoroids formed from PDX dissociated cells. We compared the dose-response of 3D tumoroids in droplets to 2D monolayer, and to tumoroids formed in microwells. Overall, this droplet platform appears as a promising tool to perform drug screening on tumoroids.

## Materials and methods

### Cell culture and chemicals

Ewing sarcoma A673 (ATCC CRL-1598) and neuroblastoma SK-N-AS (ATCC CRL-2137) cell lines were grown in DMEM (Gibco, #31966-021) supplemented with 10% (v/v) FBS (Dutscher) and 1% (v/v) penicillin/streptomycin (Gibco, #15140-122) at 37°C under 5% CO2 in humidified atmosphere. Patient-derived xenografts of Ewing sarcoma (PDX, IC-pPDX-87) were dissected, dissociated, and sorted before dilution in cell culture media (DMEM/F-12+GlutaMAX, Gibco #31331-028) supplemented with 10% (v/v) B-27 supplement (Gibco, #17504-044) and 1% (v/v) penicillin/streptomycin (Gibco, #15140-122). For tumoroids grown in droplets, cells were diluted at 500,000 cells/mL, corresponding approximately to an initial seeding of 350 cells/droplet.

For drug screening experiments, two different drugs were tested: Etoposide (10mM in DMSO, Prestwick Chemical) and Doxorubicin (10mM in DMSO, Prestwick Chemical). Both were prepared at different concentrations by consecutive dilution in the same media batch in which the cells were prepared. We also ensured that DMSO had no impact on tumoroid viability at the concentration used (Fig. S1†).

Metabolic cellular activity was determined using alamarBlue™ HS Cell Viability Reagent (ThermoFisher, #50101). For droplet experiment, alamarBlue was diluted in complete cell culture media to reach a concentration corresponding to 20% of the total final droplet volume. For experiments in wells, a volume corresponding to 10% of the media already in the wells was added to each well.

### Assessment of tumoroids growth and viability in droplet

To estimate the viability of tumoroids in droplets over a week, eight identical tubes were prepared in parallel. A “train” of droplets consisted of one droplet of cells in suspension, one droplet of culture medium and one droplet of alamarBlue. Each tube contains 20 trains.

Each day, the tumoroids of one tube were imaged, and all the 3 droplets of each train were merged. Once in contact with alamarBlue (resazurin-based assay), the viable cells of the tumoroids metabolise a fluorescent compound (resorufin) that generates a uniform fluorescent signal in all the droplet. After 12h of alamarBlue exposure, the fluorescence of each droplet was detected using a laser scanner (Typhoon FLA 9000, filter Cy3, photomultiplier value of 250V, pixel size of 25μm). The fluorescence intensity *I* was assessed by measuring the mean intensity in a rectangle centred and oriented on the droplet.

### Imaging of tumoroid

Imaging of single tumoroid was performed directly in the tubes with an inverted microscope (Nikon Eclipse Ti, magnification 10X). For equivalent diameter measurement, the area *A* of the tumoroid was measured after the segmentation of the image. The equivalent diameter *d*_*eq*_ is computed using the formula:

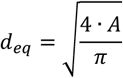

The polydispersity index (PDI) of tumoroid sizes is given by:

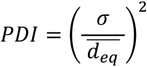

with σ the standard deviation of 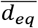

### Drug screening and IC50 determination in 2D and 3D

For drug screening on tumoroids in droplets, trains of droplets were composed of one droplet of cell suspension, one droplet of drug (at different concentration in each tube) and one droplet of alamarBlue reagent. Tubes are incubated for 24h (48h for SK-N-AS), time for the cells to aggregates and form a tumoroid. Then, a first merging is performed to add drug to the tumoroids. After 48h of drug exposure, droplets with drug-exposed tumoroids and alamarBlue reagent droplets are merged. The fluorescence detection was performed after one night of alamarBlue exposure as in the previous section. A tube without any drug was used as negative control in each experiment.The 2D drug screening assays were carried-out on 96-wells plates. 8,000 cells in 200μL were seeded per well. Plates were incubated for 24h at 37°C. Then, 50μL of drug solution was added to each well. After 48h of drug exposure, 25μL of alamarBlue was added to each. Cells without drug treatment were used as negative control, and no cells were added to blank control. Fluorescence measurement in each well was performed using a plate reader (PerkinElmer, EnSight®) with an excitation wavelength of 560nm and an emission wavelength of 590nm.Fluorescent intensities *I* were normalized as follow:

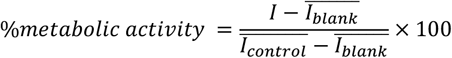

where 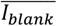 is the signal of alamarBlue without cells and 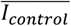 corresponds to the mean intensity of negative control (without drug).IC50 was determined using GraphPad Prism 9.3.1 by fitting the data to a 4 parameters dose-response sigmoidal curve with the least square method.^24^

### Tumoroids generation in microwells

Agarose molds (2% in PBS) were produced in a 96-wells plate with a stamp composed of 37 pillars or diameter 200μm (Fig. S2†). Cells were seeded in each well (10,000 per well), then the plate is centrifugated at 1200rpm for 4min so the cells fall into the holes. Images and fluorescence measurements were performed with a plate reader (PerkinElmer, EnSight®).

### Statistical analysis

All statistical analysis were performed using GraphPad Prism 9.3.1. After assessing the non-normality of the data, the non-parametric Mann-Whitney test was performed.

## Results

### Microfluidic setup for generating droplets in tubing

The drug screening platform consists in a XYZ stage associated to an array of 8 syringes (Cavro® XMP 6000 Multichannel Syringe Pump, Tecan Systems) (Fig. 1A). It allows to fill automatically 8 microtubes in parallel, fixed between a syringe and a travelling tube holder. The liquids are pipetted directly in the tubes from a 96-wells plate filled with the different oils, cell solutions and reagents (video of the platform in action is shown Fig. S3†). Microtubing are in polytetra-fluoroethylene (PTFE), with an internal diameter of 500μm and 50cm length (Adtech Polymer Engineering™ BIOBLOCK/12).

**Fig. 1.**
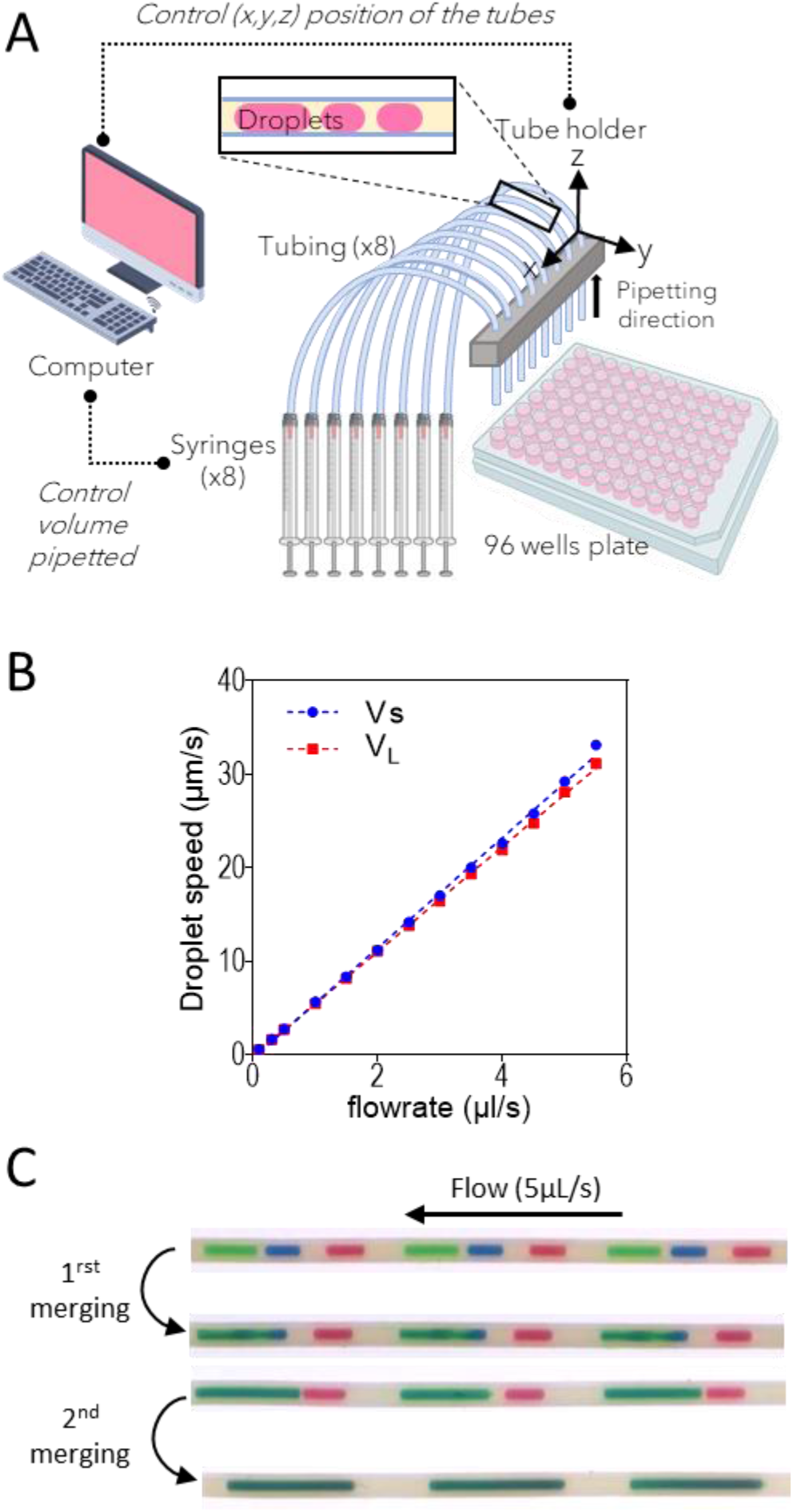
A) Set-up for generation of microdroplets in tubes. Liquids are filled in a 96-wells plate. 8 tubes of diam. 500 μm and length 50cm are fixed between syringes and a moving support that travels over the 96 wells plate in the ZY plan. Syringes pump the droplets inside the tubes. B) The mobility vs. the flowrate of the droplets for small (VS = 450nL) and big (VL = 700nL) droplets. Droplets were composed of water mixed with food colouring. C) Depending on their volume, droplets move at different speed, which allow sequential and controlled droplet merging. To merge the droplets, a flow at 5 μL/s is applied in the tube. For drug screening on tumoroids. first merging is performed after 24h incubation of the tube, between tumoroid and drug droplets. Videos of mergings are available Fig.S5 †

Each tube is filled with 20 trains of droplets (cell suspension – drug – alamarBlue). Droplets are separated by fluorinated oil with 2% surfactant (3M™ Novec™ 7500 Engineered Fluid with 2% Krytox+PEG, dSurf, Fluigent). To avoid evaporation of this oil, extremities of the tubes are filled with FC40 oil (3M fluorinert™ FC-40), separated from the surfactant by mineral oil (Sigma, M5904).Pumping is performed at 0.1μL/s for the oils and 0.2μL/s for the aqueous droplets. Tubes are reversibly clamped at both ends after filling to avoid leaks.The average volume of the droplets for cell, drug and reagent are respectively *V*_*L*_=700±50nL, *Vs*=450±55nL and *Vs*=450±55nL. In between them, the surfactant laden oil is pipetted with a volume in the range of 250-500nL. For a given pumping time, there is a small variation of droplet volume occurring due to the pressure drop variation along the length of the tube. It is compensated by an adaptation of the pumped volume following volume calibration during pipetting of droplets (Fig. S4†).

### Droplet merging

Droplet merging relies on the variation of the droplet mobility with respect to its size: larger droplets move slower than larger one,22,23 and the relative difference is flowrate dependent (Fig. 1B). This is used in the protocol to bring, on demand, two droplets of different volumes in contact to allow the merging, by pumping at a fast enough flow rate of 5μL/s here.Merging protocols (Fig. 1C) consists in applying a forward fast flow of 15μL at 5μL/s, which bring together two droplets of different sizes. Then, backward slow flow rate at 0.3μL/s is applied to bring back droplet trains at their initial position. Depending on the interdroplet spacing, these operations are repeated two to three times. The two successive merging events can fail, leading to different number of data points between the concentrations. The average success rate is between 85 and 90%.As the droplets are sterically stabilized by surfactants, there is no spontaneous merging upon contact. An electric field by ionization is applied using an antistatic gun (ZeroStat3, Milty™), following an idea proposed for demulsification process using droplet coalescence.^25^ This destabilizes the surfactant layer by charging droplet interfaces in regard with opposite charges, causing a tearing of the interface and initiating a rapid ‘unzipping’ process.Droplets are internally mixed by performing a forward and backward slow pipetting at 0.3μL/s to ensure homogeneous distributions of the molecules in the droplet.

### Tumoroid formation in the droplet platform

A single tumoroid was quasi-systematically (>99%) formed in each droplet, without any specific effort to promote aggregation. Besides, essentially all cells (>90%) are integrated into the tumoroid. The tubes are then put in the horizontal position for tumoroid maturation. Tumoroid growth has then been monitored with optical microscopy for several hours, showing cell aggregation followed by tight interactions and finally the formation of a compact spherical structure within 12 hours (videos available Fig. S6†).

### Growth and viability of tumoroids in droplets

In cell culture, the two main limiting factors for long-term culture are the gas exchanges (in particular O_2_ supply) and nutrient supply. In traditional well plate culture, gas exchanges occur quickly through the culture media/air interface and culture media can be renewed as often as needed. In the droplet platform, cells-containing droplets do not directly contact with air. PTFE and perfluoroether oil are known to be highly permeable to oxygen, among solid and liquid materials, and thus should ensure a relatively efficient gas exchange.^26^ Oxygen supply and CO_2_ removal may still be an issue, which had to be checked. Also, in the workflow proposed here, the medium is not renewed for the first 3 days.To check for the above potential problems, the growth and the viability of tumoroids were investigated for tumoroids made from A673 cells (Ewing sarcoma). Tumoroids were grown in 700nL droplets for several days. Each day, the size of each tumoroid and its metabolic activity were assessed. After 1 day of maturation, tumoroids measure 122±21 μm of diameter (PDI=0.03). They growth and reach 192±13μm (PDI = 0.005) after 4 days. The metabolic activity of the tumoroids increases for the first 3∼4 days before reaching a plateau and slightly decreasing (Fig. 2A-B). The increase in metabolic activity is proportional to the tumoroid growth (Fig. 2C), which diameter increases over 3 days before saturating.

**Fig. 2.**
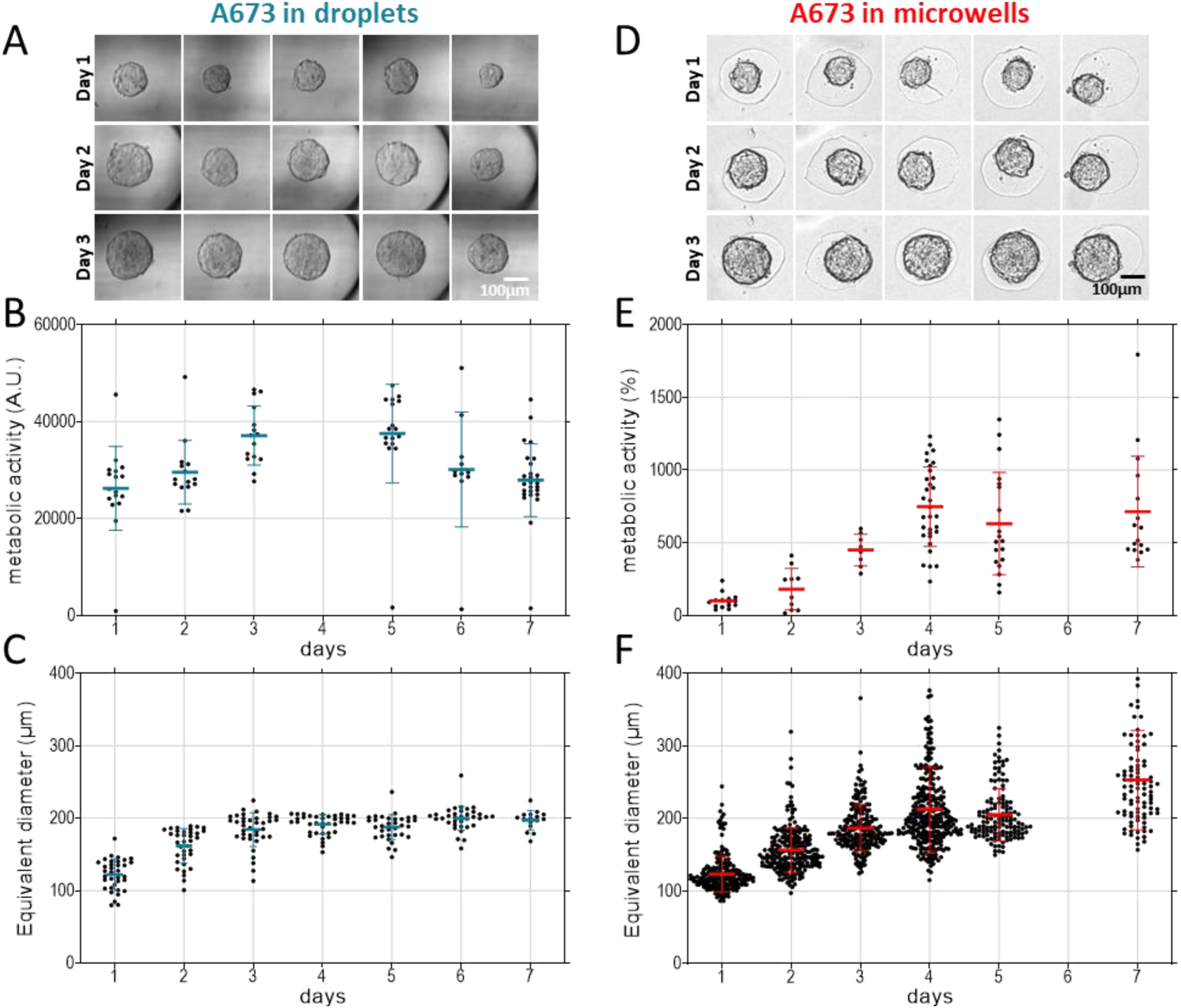
Growth of tumoroids (A673) in droplet and microwell platforms. A) Images of tumoroids, B) evolution of metabolic activity and C) equivalent diameter of tumoroids grown in droplets. 350 cells were seeded in 700nL droplet. D) Images, E) evolution of metabolic activity and F) equivalent diameter of tumoroids grown in agarose microwells. One point corresponds to one tumoroid, except for metabolic activity in microwells (where it corresponds to one well of 37 tumoroids). Errors bars represent SD of the mean.

To further check if the growth of tumoroids in droplets could be possible but sub-optimal, tumoroids were also prepared in agarose microwells,^27–29^ starting from the same number of cells. The diameter of the holes has been fixed at 200 μm to match the final size of tumoroids in droplets. The metabolic activity and the diameter of the tumoroids were measured every day over a week. Figure 2D shows a panel of the tumoroid growth. Remarkably, tumoroid growth in droplets and in microwells are very similar, with tumoroids metabolic activity (Fig. 2E) and diameter (Fig. 2F) also increasing in microwells over the 3 to 4 days before reaching a plateau.Our droplet platform appears then as a robust tool to generate reproducible and monodisperse tumoroids during a period of at least 4 days, with extremely rare failures observed during the experiments (<1%). The platform is thus well adapted to the drug testing workflow, in which drug exposure is started after 1 days of tumoroid maturation, and the metabolic activity is checked at day 3, which is still in the growth regime in the absence of drug.

### Drug screening on tumoroids

Drug screening was then carried out by establishing dose-response curves to chemotherapy. Figure 3 focuses on the exposure to the chemotherapeutic drug etoposide, and compare the drug-response for A673 cells grown as two-dimensional (2D) monolayers (Fig. 3A-B), or as tumoroids in droplets and in microwells (Fig. 3C-D). The curves are normalized with respect to the no-drug case and after subtracting the residual signal with only alamarBlue (without drug and cells: this residual signal is due to the initial baseline fluorescence of the substrate). The half maximal inhibitory concentration (IC50) of these curves have been determined (Table 1).

**Table 1.** IC50 values. Etop: Etoposide. Dox : Doxorubicin. * spheroid in microwells

| Cell type | Drug | Spheroid in droplet |  | Monolayer |  |
| --- | --- | --- | --- | --- | --- |
|  |  | IC<sub>50</sub> (μM) | 95%CI | IC<sub>50</sub> (μM) | 95%CI |
| A673 | Etop | 5.9 | [5.3 ; 6.7] | 3.0 | [2.3 ; 3.9] |
|  | Etop* | 4.1 | [2.3 ; 12.1] |  |  |
|  | Dox | 1.4 | [1.1 ; 1.7] | 0.4 | [0.4 ; 0.5] |
| SK-N-AS | Etop | - |  | - |  |
|  | Dox | 3.0 | [1.9 ; 7.3] | 1.0 | [0.6 ; 2.1] |
| PDX | Etop | 3.4 | [2.6 ; 4.7] | 0.5 | [0.5 ; 0.6] |
|  | Dox | 1.2 | [ - ; 1.7] | 0.2 | [0.2 ; - ] |

**Fig. 3.**
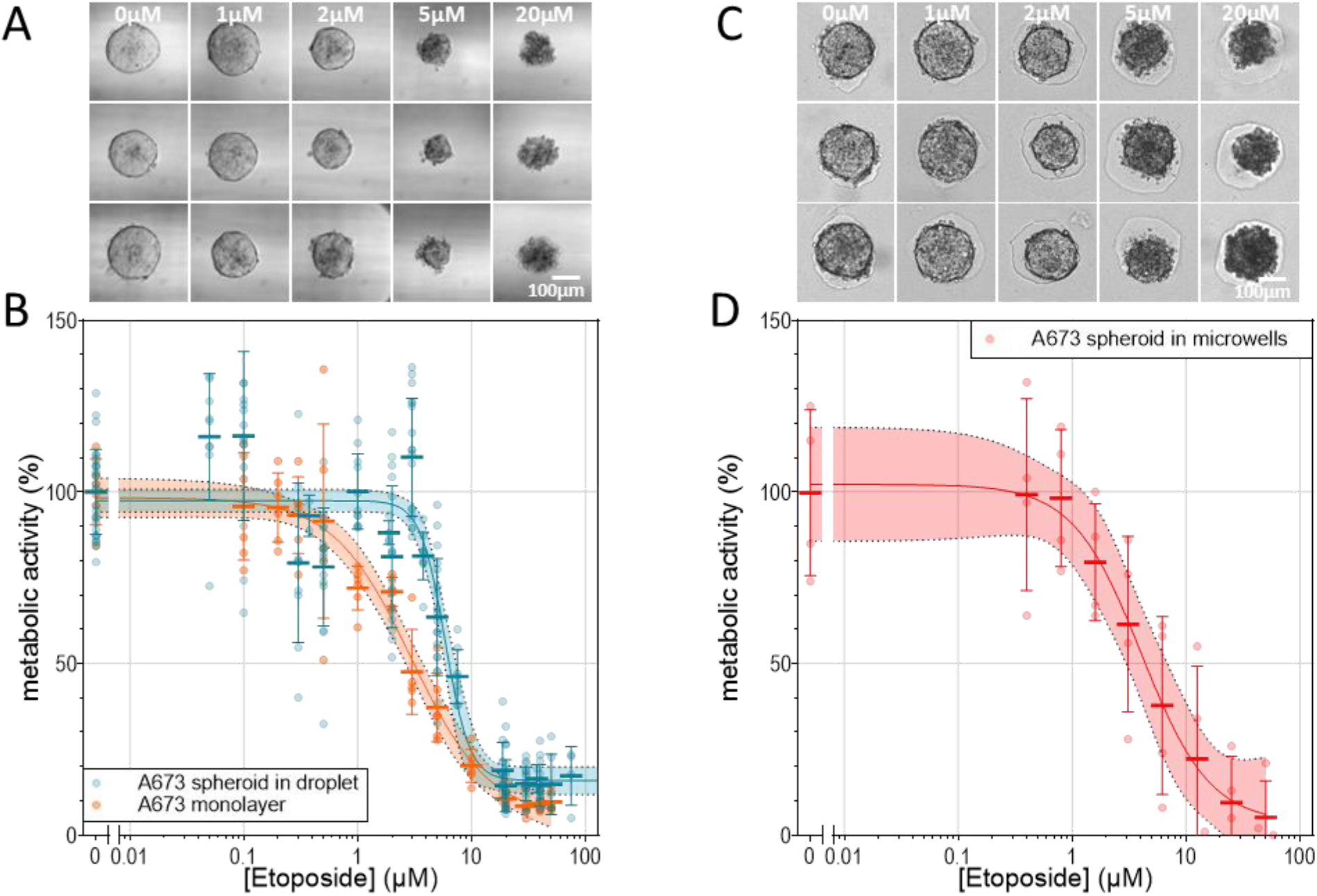
Drug screening of A673 tumoroids with etoposide. A) Images of A673 tumoroids in droplets exposed to different drug concentrations. B) Dose-response curve of A673 tumoroids and monolayer. C) Images of A673 tumoroids in microwells exposed to different drug concentrations. D) Dose-response curve of tumoroids in agarose microwells. Cells were seeded for 24h and exposed to drug for 48h. Metabolic activity was determined using alamarBlue assay. Initial cell number was about 8,000 cells per well in monolayer and 350 cells per droplet in tumoroids. Each point represents one tumoroid or microwell. Errors bars represent SD on the mean. Straight line represents the curve fitting to a 4-parameter sigmoid, with its 95% Cl.

A second chemotherapeutic compound, doxorubicin, was also tested in the droplet platform on A673 tumoroids (Fig. 4A), and compared again with the response in 2D configuration. As for etoposide, the drug screening performed well at single Spheroid in droplet Monolayer tumoroid level, with the correct determination of IC50. A second cancer cell line, neuroblastoma (SK-N-AS), was further tested in the same settings, upon administration of etoposide (Fig. 4B) and doxorubicin (Fig. 4C). The dose-response curve was relevant for doxorubicin exposure, with accurate IC50s determination in both 2D and in tumoroids in droplets, as indicated in Table 1. By contrast, the SK-N-AS cell line appears to be low-responsive to etoposide, both in 2D and in droplets, which is consistent with the high IC50 values found in the literature.^30–32^ Thus, IC50 could not be inferred but it validates that the droplet platform can also be indicative of an absence of efficiency of a given drug. Overall, IC50 values obtained here in 2D monolayers are close to the ones reported in the literature for the same cells and drugs (Table S1†). Interestingly, in 3D tumoroids grown in droplets, the IC50 is higher than for cells in 2D configuration. A systematic statistical analysis was performed to compare metabolic activity at the same drug concentration between 2D and 3D. A significant difference (p < 0.001) was systematically found around the IC50 concentrations (Fig. S7†).

**Fig. 4.**
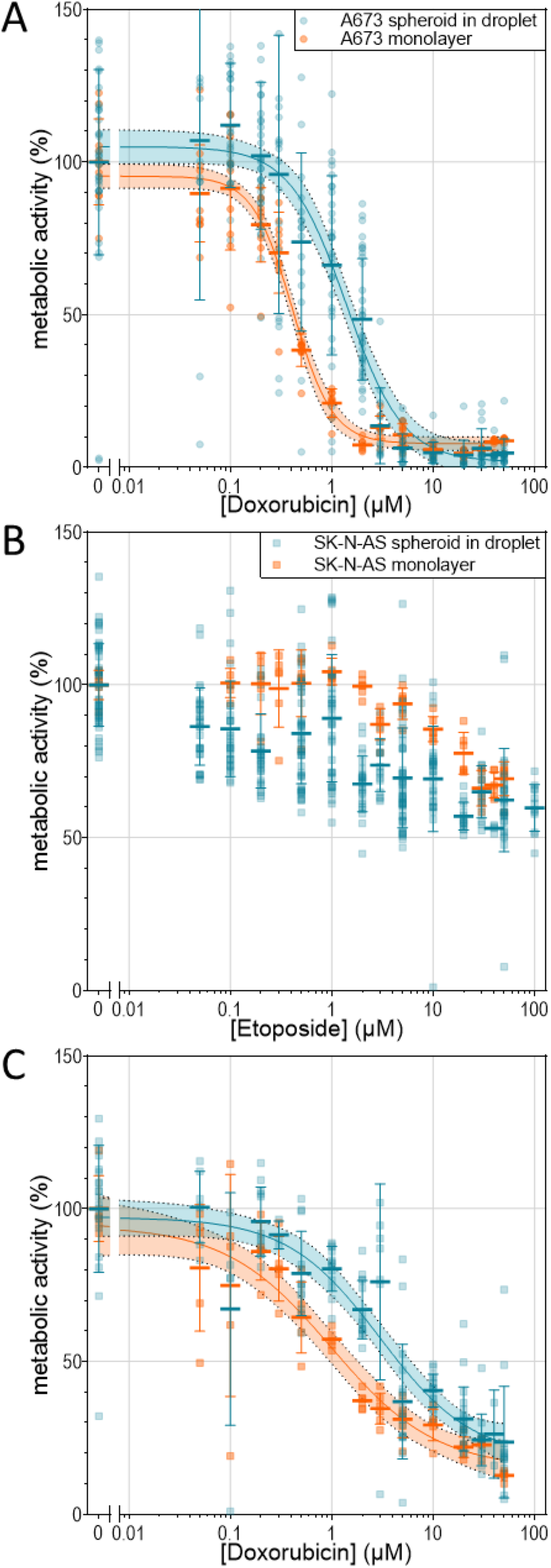
Dose response curves of A) A673 exposed to doxorubicin, B) SK-N-AS exposed to etoposide and C) SK-N-AS exposed to doxorubicin, in droplets tumoroids and in monolayer. A673 were seeded for 24h and SK-N-AS for 48h, then exposed to drug for 48h. Metabolic activity was determined using alamarBlue assay. Initial cell number was about 8,000 cells per well in monolayer and 350 cells per droplet in tumoroids. Each point represents one tumoroid or well. Errors bars represent SD on the mean. Straight line represents the curve fitting to a 4-parameter sigmoid, with 95% Cl.

The endpoint of the platform is to be used on patient cells in a personalized medicine perspective. To prepare evolution towards this clinical situation involving primary cells, drug screening was finally performed on, PDX derived from Ewing’s sarcoma (similar to A673). PDX is expected to be closer to primary patient cells than cell lines, and was thus considered here a good model to address the applicability of the platform to primary cells. A first observation is that tumoroids could be successfully grown from PDX in the droplets platform. They were then exposed to etoposide (Fig. 5A for typical images, and 5B for the curve) and doxorubicin (Fig. 5C), and a similar drug testing was performed in 2D for comparison. A second observation is that a larger variability in the response of the tumoroids than for cell lines is observed, both for spheroids in droplets and monolayers, but one can clearly see an effect of the drugs on the cells and IC50 determinations could be achieved with statistical significance Table 1. Here, the difference between the 2D monolayer configuration and the tumoroid one is even more important than for the cell lines.

**Fig. 5.**
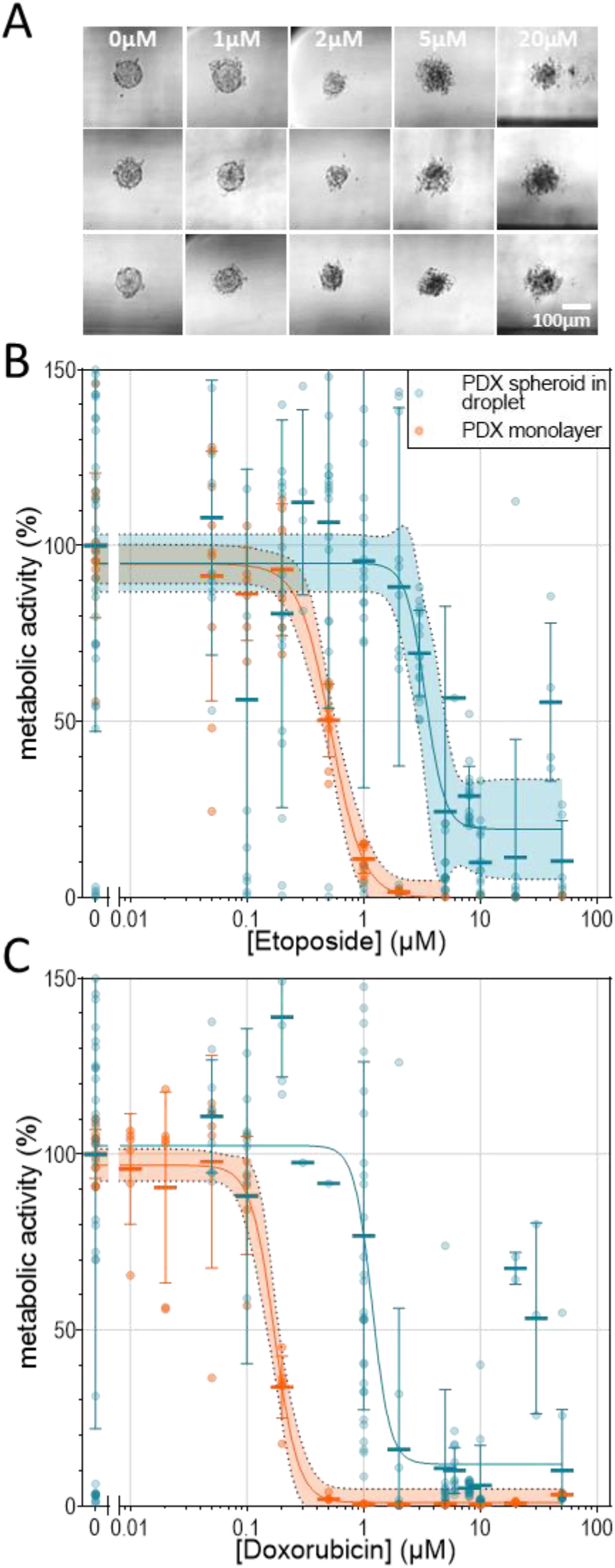
Drug screening on PDX. A) Images of PDX tumoroids in droplets after drug exposure. B) Dose response curve of PDX tumoroids exposed to etoposide. C) Dose response curve of PDX tumoroids exposed to doxorubucin. Cells were seeded for 24h and exposed to drug for 48h. Metabolic activity was determined using alamarBlue assay. Initial cell number was about 8,000 cells per well in monolayer and 350 cells per droplet in tumoroids. Each point represents one tumoroid or well. Errors bars represent SD on the mean. Straight line represents the curve fitting to a 4-parameter sigmoid, with 95% Cl. For C), 95% Cl was too wild and does not appear.

## Discussion

The platform proposed here combines the advantages of growing three-dimensional tumoroids in sub-microliter droplets, to reduce cells consumption, and of droplet merging on demand to submit tumoroid to drugs and assess their impact through metabolic assay. Each droplet thus serves as a micro-reactor platform by providing a non-adherent environment, and allowing the implementation of complex protocols implying the addition of different reagents at preselected times, and high-resolution imaging.

### Technical issues and possible evolutions

#### Pipetting accuracy and reproducibility

Once corrected from the systematic bias due to pressure drop, which depends essentially on the tube diameter and pipetting speed, and can be well corrected thanks to a pre-calibration, the uncertainty in droplet volume is about 50nl, i.e., for the volumes chosen here, of the order of 10% at most. 10% is also the order of magnitude of the initial fluctuation in the number of the pipetted number of cells, even in a perfectly known volume, and we consider that a 10% error in drug concentration is acceptable for the performance of IC curves, considering the number and range of concentrations used, and the observed intrinsic variation in cells response.

#### Number of droplet trains in a tube

The number of droplet trains in a tube is essentially limited by the length of the tube. As discussed, the pressure drop can be taken into account, and the number of 20 trains, used in this study, is not a limit in this respect. However, the handling of very long tubes can be tricky, in particular for imaging. We thus consider that 20 trains is a reasonable compromise, and that increasing the number of tubes used in parallel, beyond the value of 8 used here, would be a better way to increase throughput.

#### Imaging

Imaging is performed in a planar configuration, and can be achieved either in a microscope (preferably inverted) or in plate readers (the latter possibly requiring an adaptation, depending on the details of the instrument). Observation is performed in-tube, which involves some distortion close to the tube walls, due to refraction. This distortion is minimized by “sandwitching” the tubes in fluorinated oil between planar glass slides, and by the fact that spheroids spontaneously position along the droplet’s axis, where imaging distortion is minimal. The tube thickness provides another limitation, imposing relatively long working distance (typically > 2mm) and low magnification (typically up to 10x).

#### Droplet merging

As presented above, droplet merging is performed by asymmetric “push-pull” pumping at different speeds. This allows to impose freely the timing of different merging events, in theory up to an arbitrary number of droplets, by playing on the distance between different droplets. The fluctuations in droplet sizes, and time considerations, will in practice limit the conveniently achievable number of merging events, but one can go significantly beyond the number of 2 merging events used here, if needed. Due to the rare but possible presence of air bubbles and variations in the oil volume between droplets, the merging steps can fail for some trains on droplet. The average success rate has evolved during the study, due to progressive optimization of the interdroplets spacing. Initially around 60%, it is now around 85∼90% in our laboratory prototype, and has room for improvement during maturation of the method and industrialization of the instrument and protocols, towards a success rate of at least 95%.

#### Throughput and hands-on time consideration

The longest steps in the workflow, by far, are the incubation ones, so we chose to dissociate the pipetting from the incubation steps: this way the platform for pipetting can be used continuously, and incubation can be performed in conventional incubators. We also chose to perform imaging in conventional microscopes or scanners, to benefit fully from the high technological maturity and scales economy of existing imaging platforms. The innovation in the method thus lies in the pipetting platform, which can be used in continuous to prepare removable casing holding an array of tubes, with a spacing equal to the spacing of vials in a MTP. At present, we close tubes before removal from the platform by manual clamping, and in routine this imposes a typical hands-on time of 10-15 min every hour for tube array exchange. In our present prototype, which was not optimized for throughput, the total occupation time of the pipetting machine for a full workflow (including initial pipetting and two merging events) is around 1h30 hour per tube array (8 tubes in parallel, i.e. 160 droplet trains). The current potential throughput is around 1000 data points a day for a 9-hours per day working schedule.In the specific application of drug screening by IC50 tested here, considering that, according to our results, 20 replicates provide statistically significant results, and an IC50 typically requires 10 different concentrations, this would represent a throughput around 5 IC50 curves with 20 replicated data point per day. This is definitely lower than what can be currently achieved in 2D screening in 384 wells plates, but of course, transposition of the technology to routine diagnosis would imply multiplication of the number of tubes operated in parallel, and overnight automation of tube array transfer with a robotized platform, as already implemented for MTP.

### Biological issues and perspectives

#### Mechanisms for spheroid formation

We noticed in the results section that individual spheroids form with a high yield and consistency in each droplet. Such remarkable tumoroid formation may have several explanations. During the tubes filling process, all pre-existing droplets move along the tube, generating convective recirculation inside the droplets: the strong shear occurring inside the lubrication oil film between the droplet and the tube wall induces, in the cylindrical section of the droplets, a tubular flow of the droplet’s content with regards to its center of mass. By mass conservation, this flow is compensated by a flow in the opposite direction along the droplet’s axis. This recirculation creates two stagnation zones in the vicinity of the two droplet apexes, towards which all cells contained in the droplet are dragged, creating a local concentration of cells promoting aggregation.^33^ Symmetry breaking between the two apexes, leading to a single aggregate instead of two, is probably induced by gravity and cell sedimentation, since the tube is maintained in vertical position during the pipetting. This effect can be maximized by keeping the tubes in a vertical position for 15 minutes after the end of pipetting. This process is indeed reminiscent of the hanging drop approach, except that it is accelerated by intra-drop convection.

#### Growth and viability of spheroids

Experimentally, the growth of spheroids during the 4 first days is unaffected by confinement and nutriment or oxygen shortage, as shown by comparison with open microwells (see Results section). This is sufficient for many applications, and notably the drug screening performed in this article, but growth during longer periods could be of interest in other applications. We did not investigate here specifically the cause of growth saturation after 5 days. The problem of nutriments shortage (and to some extent toxic waste elimination) could be easily solved by adding to the protocol additional merging events with “fresh” medium droplets inserted in the train of drops. The problem of oxygen supply may be provided through changes of materials and/or geometry. Finally, the size of droplets is also a tunable parameter, and it may be expanded to volumes of several microliters by a simple change in the tube diameter.

#### Difference in IC50 between platforms

The comparison with conventional 2D culture showed a systematic increase in IC50 for 3D culture as compared to 2D, both in droplets and in microwells. This highlights the lower sensitivity of cancer cells to drugs in 3D configuration compared to 2D, logically reflecting the difference in drug penetrability within the tumoroid. Comparison between tumoroids grown in droplets and microwells yielded very similar IC50’s confirming that the difference with conventional 2D assays is mainly due to the 3D nature of the cell’s organization, and not to the droplet platform. Note, however, that in contrast to microwells, in which tenths of tumoroids are grown in a single well, the droplet platform performs screening at the single spheroid level, which has pros and cons, as discussed below with respect to PDX.

#### The case of PDX

PDX experiments systematically yield more dispersion in results, both in 3D and in 2D. We attribute this to an expected higher intratumoral heterogeneity, since the constitution of cell lines involves a selective evolution that is expected to be less pronounced in PDX (indeed one of the main reasons why the latter are considered more physiological). Although our experiments on PDX are at this stage too limited to jump to definite conclusions, we note that the dispersion is higher for the droplet platform than for the 2D assay. This may be due to the fact that this platform works at the single spheroid level, combined with the small number of cells used for each spheroid. This creates additional noise (compensated by the systematic use of 20 replicates, which could be increased if needed), but on the positive side, it may provide a new tool to address intratumoral heterogeneity. This will probably require more elaborate assays beyond the philosophy of this first proof-of-concept experiment, but we nevertheless expect this to be an interesting potential of the platform. Along this line, too, one should note that since individual spheroids are deterministically identified by their rank in the tube, they can be pipetted back and collected individually for further analysis, e.g. by next-generation sequencing (NGS). There is only single tumoroid inside each droplet, so that drug testing directly correlates with a single unit. The dispersion in our results, as compared to 2D, suggests that even at the level of a few hundreds of initial seeded cells, heterogeneity between samplings can play a role, and different tumoroids can have a different resistance to the drug.

#### Number of cells per assay

In our view, a major advantage of the platform is the low number of cells needed for each assay. Here, we typically used 350 initial cells per droplet, and this may not be ultimate lower limit, although we did not at this stage investigate systematically the effect of cells content on the results robustness. Considering that, according to our results, 20 replicates provide statistically significant results, and an IC50 typically requires 10 different concentrations, around 70,000 cells are required for a complete IC50 assay. To our knowledge, this is at least 10 times less than with conventional methods. A systematic study by Rajer *et al*.^15^ showed that a Fine Needle Aspiration, a very common and weakly invasive biopsy method, provides in average between 1 and 5×10^6^ of cells (more specifically, in their hands, the fraction of FNA with less than 5×10^5^ cells is 4.9% for lymph nodes, 16.7% for breast and 18% for thyroid). Thus, with this platform, a single FNA should provide in most cases the possibility to test at least 10 different drugs or drug combinations. We thus believe that it opens the route to direct personalized drug screening on primary cells from patients.

#### Comparison with other droplet platforms

To our knowledge, the two closest systems to this droplet microfluidics platform come from the groups of Merten^20^ and Baroud.^21^ In the first system, 100nL droplets in which cells, drug and caspase assay are encapsulated are prepared in continuous. The liquids are injected in the channels through Braille valve to prepare the cocktails of drugs coupled with dyes and cells. This system can be used to test up to 10 drugs and their combinations (56 conditions in total) on 12 replicates for each condition. This system is a powerful tool to test lot of different drug combinations, but it requires a microfabricated chip and valves system to fill the channels, as well as a barcoding system to read the results. Also, drugs are tested on cells in suspension, and not on 3D cells organoids. The prototype proposed has a significantly higher throughput than the one proposed in the present article, but, importantly, it does not allow droplet merging, and thus it is much more limited in terms of protocols flexibility. This is indeed why the assay proposed was a caspase assay performed directly on cell suspensions. This assay is specific to apoptosis, and it is thus not well adapted for the screening of drugs with modes of action not directly involving apoptosis. The second system involves trapping cells into droplets anchored on a microfluidic chip in “capillary traps”, and form tumoroids (up to 252 tumoroids per chip). One additional droplet can be merged on demand to this first one, containing other cells types for co-culture assays, or drugs for drug screening experiments. The authors managed to test 26 compounds on one chip, detected with a bar coding system.As compared to these systems, our platform thus has the following advantages: it does not require barcoding, since the “identity” of sample and reagent droplets is built in the invariable droplet “rank” in the tube. It also allows the implementation of complex protocols, involving several sequential mergings of droplets. Finally, since it works at the single spheroid level, the consumption of cells is minimal, which can be highly precious for clinical or rare samples. These advantages are currently paid by a lower throughput, but the latter could be strongly improved by parallelization and automation.

## Conclusions

A droplet microfluidic system to miniaturize drug screening assays on organoids is introduced, based on a purely hydrodynamic framework. This system can be easily implemented with minimal infrastructure and material. The miniaturization of the process results in a reduction of compounds, reagents, and most importantly of tumour cells. Tumor spheroids (from cell lines and PDX) were grown in droplets from aliquots containing 350 cells in average, in sub-microliter droplets for several days. This platform can reliably generate the IC50 curve from drug screening experiments, using a total of 70,000 cells for a full IC50 curve and 20 replicated experiments. The possibility to grow tumoroids from only few hundred cells makes this system suitable for drug testing on primary cells, generally available in fewer quantity than provided by cells amplification or PDX, and open the way to direct high-throughput assays for treatment orientation. Ultimately, it could be used to test combinatory therapies of drugs added sequentially to the tumoroids. Finally, and importantly, the system developed here could be a highly valuable asset to dissect the mode of actions of drugs by using several biological markers on the same tumoroids to identify the different cell death mechanisms involved, since it allows multiple merging events with droplets containing different reagents. It thus appears that the potential of the droplet platform for tumoroids culture could have used for both treatment development and for cancer precision and personalized medicine.

## Supporting information

ESI

## Author Contributions

OD, DS and JLV conceptualized the research. OD, CW and JLV participated to funding acquisition. YZ developed the prototype and instrument management software and performed preliminary experiments. VA implemented evolutions of the instrument. DS and OD defined biological and clinical demand and provided PDX samples. CP and KRM developed experimental protocols and performed experiments. CW and CP performed control experiments in microwells. CP, KRM and CW developed imaging protocols and data analysis methodology, and analyzed data. CW and JLV supervised the research. CP, CW, KRM and JLV wrote the original draft. All coauthors participated to reviewing and editing.

## Acknowledgements

We are indebted to Sakina Zaidi for the preparation and provision of cells from PDX, and to Dr Julien Autebert for major suggestions for the implementation of droplets merging. This work was supported in part by ERC-POC COMMiT (ERC-2019-PoC-899537) and by ANR PRCE DROMOS (<u>Project-ANR-20-CE19-0012</u>); CP acknowledge a PhD fellowship PSL-Qlife ; YZ acknowledge a fellowship from ARC Fondation pour la Recherche sur le Cancer, #PDF20180507309. Fig. 1A was created with the help of BioRender.com.

