## Supplementary material for "Simple droplet microfluidics platform for drug screening on cancer spheroids": ESI

### Electronic Supplementary Informations

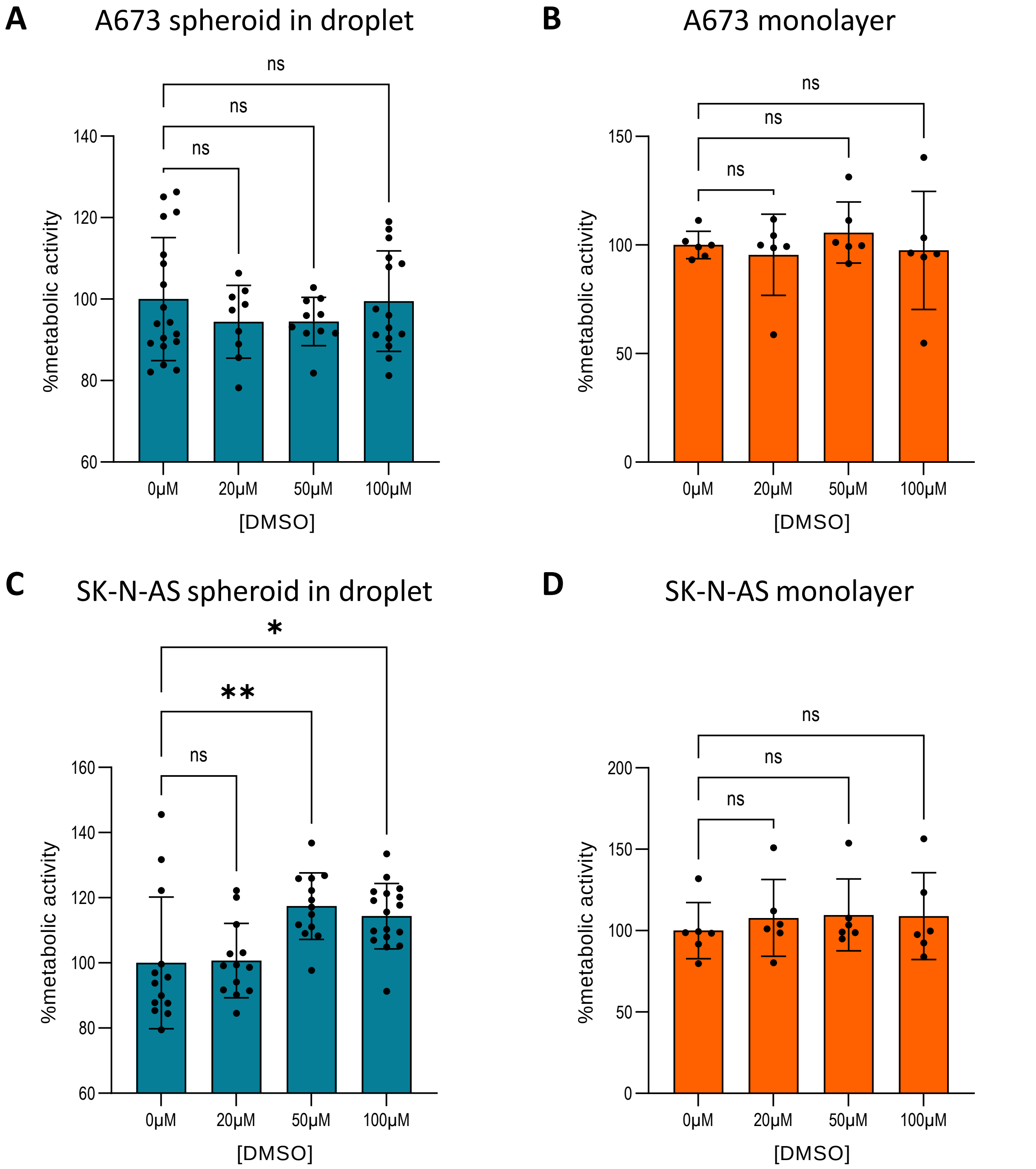


**Fig. S1** Impact of DMSO on cells at different concentration for A) A673 spheroid in droplet, B) A673 monolayer, C) SK-N-AS spheroid in droplet and D) SK-N-AS monolayer. Conditions were similar as ones used with drug: cells were seeded for 24h and exposed to DMSO for 48h. Metabolic activity was determined using alamarBlue assay. Initial cell number was about 8,000 cells per well in monolayer and 350 cells per droplet in spheroid. Each point represents one tumoroid or well. Errors bars represent SD on the mean. Data did not pass normality test, non-parametric Kruskal-Wallis test was performed using GraphPad Prism 9.3.1. *** $p < 0.001$, ** $p < 0.01$, * $p < 0.05$.

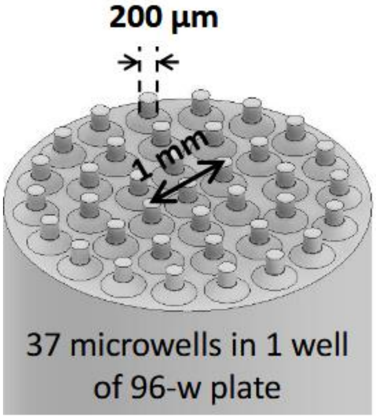


**Fig. S2** Scheme of the stamp for agarose microwells production. 34 pillars of diameter 200µm are distributed over a cylinder fitting in a well of a 96-wells plate.

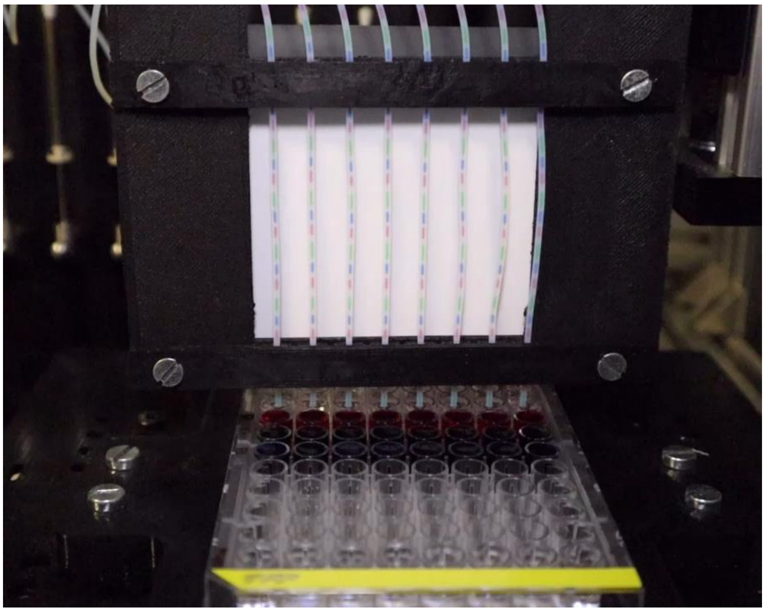

**Fig. S3 (Caption)** Video of the drug screening platform during filling of 8 tubes with colored droplets (food coloring in water). Real speed.

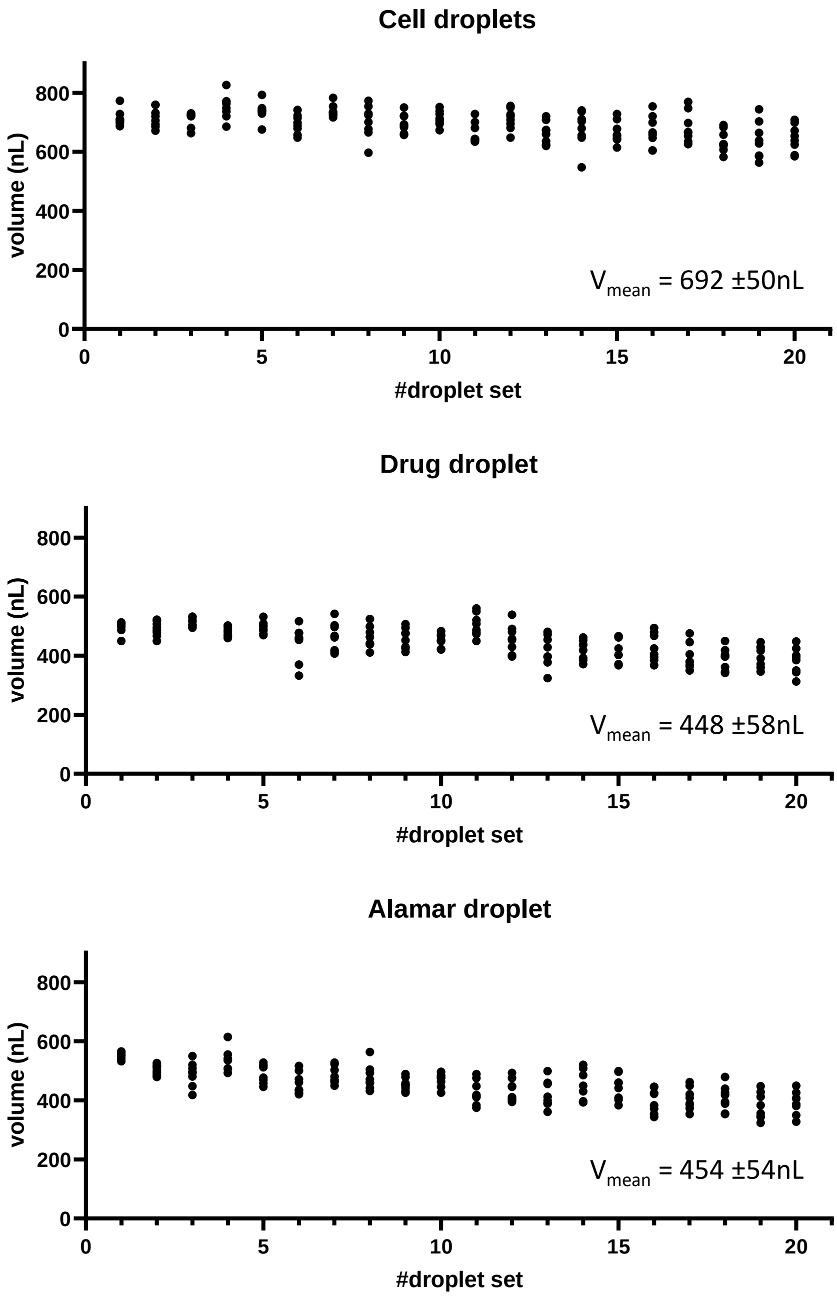


**Fig. S4** Volumes of cell, drug and alamar droplets. Droplets were generated by pipetting 20 trains of colored droplets in each of the 8 tubes in parallel. Volumes were computed from the length of the droplets measured on fluorescence images of the tubes.

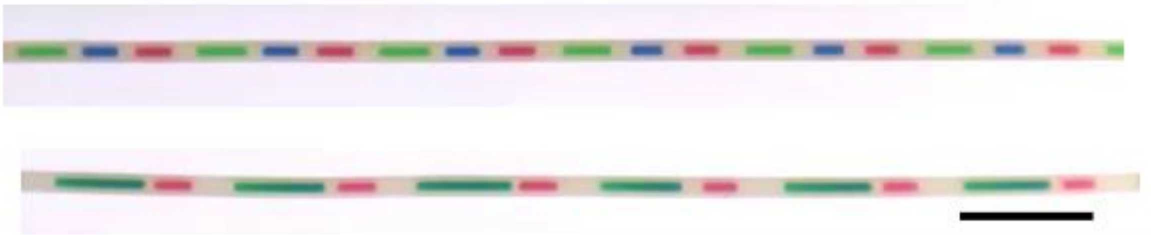

**Fig. S5 (Caption)** Video of the mergings of droplets. Top shows the first merging, bottom the second. Droplets were made of food coloring diluted in water. A fast flow (5µL/s) is applied, then a slow flow (0.3µL/s) to bring the droplets back to their initial position. This operation is repeated twice for each merging. Speed x10. Scale bar: 1cm.

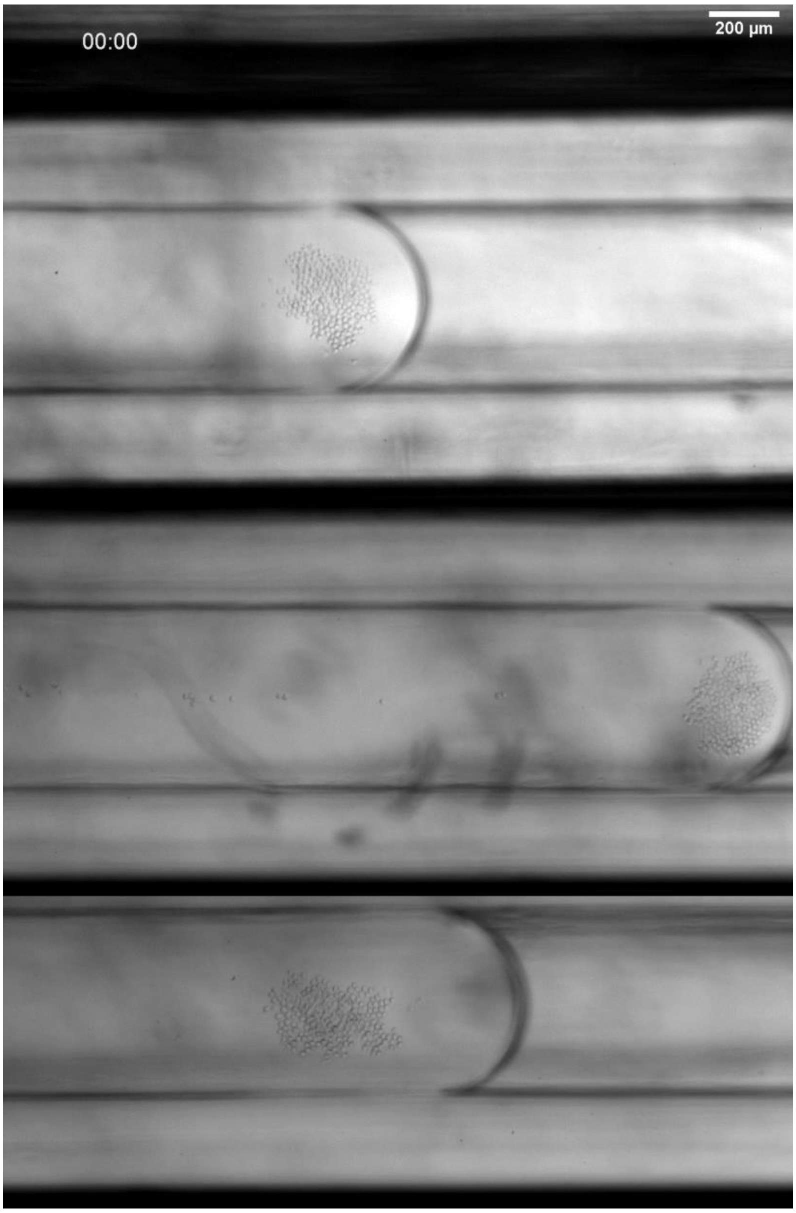


**Fig. S6 (Caption)** Video of tumoroid formation for 17h. Images were made every 1min for 4h, then every 20min in microscope (Nikon Ti, magnification 4x) with a controlled environment chamber (5%$CO_2$, 37°C).

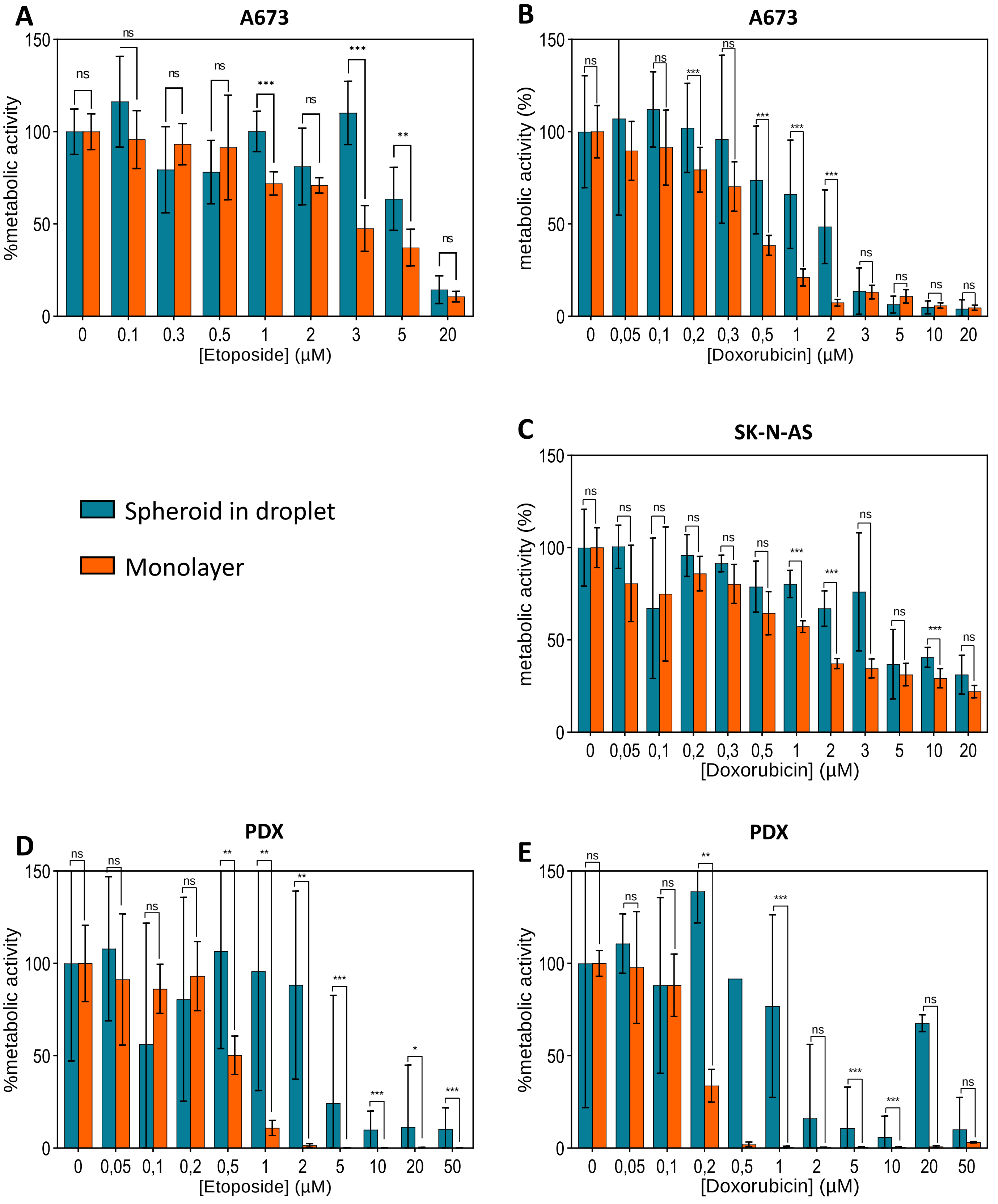


**Fig. S7** Comparison of the impact of drug concentration on metabolic activity between spheroids in droplet and monolayer culture. Metabolic activity was determined using alamarBlue assay. Initial cell number was about 8,000 cells per well in monolayer and 350 cells per droplet in spheroid. Errors bars represent SD on the mean. Mann-Whitney test was performed using GraphPad Prism 9.3.1. *** $p < 0.001$, ** $p < 0.01$, * $p < 0.05$.

| Cell line | Drug | Exposure time | Experiment design | Assay | IC50 | Ref |
|---|---|---|---|---|---|---|
| A673 | Etoposide | 96h | 0.5-1 $10^4$ cells/well, 96 wells plate | MTS | 0,88µM | Boehme *et al.*[1] |
| | | 24h | 200,000 cells/well, 6 wells plate | Flow cytometry (7-AAD & Annexin V-FITC) | >200µg/mL = >340µM | Chevalier *et al.*[2] |
| | Doxorubicin | 96h | 0.5-1 $10^4$ cells/well, 96 wells plate | MTS | 27,18nM = 0,027µM | Boehme *et al.*[1] |
| | | 24h | 200,000 cells/well, 6 wells plate | Flow cytometry (7-AAD & Annexin V-FITC) | 2µg/mL = 3,7µM | Chevalier *et al.* [2] |
| SK-N-AS | Etoposide | 48h | 3 000 cells/well, 96 wells plate | MTT | 80µM | Das *et al.*[3] |
| | | 48h | 1 $10^4$ cells/well 96 wells plate | MTS | Not determined but similar, only 4 points | Day *et al.*[4] |
| | | 7 days | 1 $10^3$ cells/well 6 wells plate | Flow cytometer (Caspase-GloTM3/7) | 0,09µg/mL = 0,15µM | Harvey *et al.*[5] |

**Tab. S1** Some IC50 values from the literature.
